# Pregnancy-associated plasma protein-aa promotes neuron survival by regulating mitochondrial function

**DOI:** 10.1101/456632

**Authors:** Mroj Alassaf, Emily Daykin, Marc Wolman

**Affiliations:** Department of Integrative Biology. University of Wisconsin, Madison, Wisconsin, United States of America; Neuroscience Training Program. University of Wisconsin, Madison, Wisconsin, United States of America

## Abstract

A neuron’s longevity is regulated by both extracellular molecular factors and the regulation of its intracellular functions, including mitochondrial activity. It remains poorly understood which extracellular factors promote neuron survival by influencing mitochondrial function. Through zebrafish mutant analysis, we reveal a novel extracellular neuronal survival factor: Pregnancy-associated plasma protein-aa (Pappaa). Neurons in *pappaa* mutant larvae die precociously and exhibit multiple mitochondrial defects, including elevated mitochondrial calcium, membrane potential, and reactive oxygen species production (ROS). In *pappaa* mutants, neuron loss is exacerbated by stimulation of mitochondrial calcium load or ROS production and suppressed by exposure to a mitochondrial ROS scavenger. As a secreted metalloprotease, Pappaa stimulates local insulin-like growth factor 1 (IGF1) signaling; a known regulator of mitochondrial function and neuron survival. In *pappaa* mutants, neurons show reduced IGF1-receptor activity and neuron loss is attenuated by stimulation of IGF1 signaling. These results suggest Pappaa-IGF1 signaling promotes neuron survival by regulating mitochondrial function.

## Introduction

Without a sufficient regenerative capacity, a nervous system’s form and function critically depends on molecular and cellular mechanisms that promote neuron longevity. A neuron’s survival is challenged by its own energy demands. Considerable energy is required for basic neuron functions, including maintaining membrane potential, propagating electrical signals, and coordinating the release and uptake of neurotransmitters (Halliwell, 2006; Kann and Kovács, 2007; Howarth et al., 2012). A neuron’s metabolic energy is primarily supplied by mitochondrial oxidative phosphorylation, a process in which the flow of electrons across the electron transport chain produces adenosine triphosphate (ATP) (Kann and Kovács, 2007). Although this process is essential to neuron survival, a consequence of mitochondrial oxidative phosphorylation is the generation of cytotoxic reactive oxygen species (ROS). The oxidative stress caused by ROS accumulation damages vital cell components including DNA, proteins, and lipids (Schieber and Chandel, 2014). Neurons are particularly vulnerable to oxidative stress due not only to their energy needs and thereby ROS production, but also to their relatively insufficient antioxidant capacity compared to other cell types (Halliwell, 1992). Cumulative oxidative stress can yield neuron loss, as observed in aging and neurodegenerative disorders including Alzheimer’s disease (AD), Parkinson’s disease (PD), and Amyotrophic lateral sclerosis (ALS) (Perry et al., 2002; Barber et al., 2006; Mattson and Magnus, 2006; Blesa et al., 2015). Thus, regulation of mitochondrial ROS production and a neuron’s capacity to minimize oxidative stress, are critical determinants of neuron survival.

The insulin-like growth factor-1 (IGF1) signaling pathway is known to affect a neuron’s mitochondrial function and its survival. Inhibition of IGF1 signaling causes loss of hippocampal neurons (Luo et al., 2003) and reduced IGF1 signaling has been shown to disrupt mitochondrial function, biogenesis, and ROS production (Lyons et al., 2017). Such mitochondrial defects are detrimental to neuron survival (Schon and Manfredi, 2003; Golpich et al., 2017). Coincident with age-related neuron loss, IGF1 levels decrease (Hammerman, 1987). In humans, low IGF1 levels show comorbidity with brain atrophy and dementia in AD, whereas high IGF1 levels are associated with decreased risk of AD dementia and greater brain volume (Westwood et al., 2014). In rat models of aging, old rats show reduced IGF1 and exhibit mitochondrial dysfunction, oxidative damage, and increased expression of pro-apoptotic genes in the brain. Exogenous IGF1 treatment reverses these outcomes (García-Fernández et al., 2008) and has also been demonstrated to protect motor neurons and delay symptomatic progression in a mouse model of ALS (Sakowski et al., 2009). Combined, these findings implicate IGF1 signaling in supporting neuron survival by regulating mitochondrial function and suggest that modulating IGF1 signaling has therapeutic potential for neurodegenerative disease.

It remains poorly understood how endogenous IGF1 signaling is regulated to influence a neuron’s survival and mitochondrial activity. IGF1 is synthesized both in the liver for systemic distribution and locally in tissues, including the nervous system (Bondy et al., 1992; Sjögren et al., 1999). IGF1’s biological functions are mediated by binding to cell membrane bound IGF1 receptors (IGF1Rs), which act as receptor tyrosine kinases. When bound by IGF1, the IGF1R autophosphorylates and stimulates intracellular PI3kinase-Akt signaling (Feldman et al., 1997). Extracellularly, IGF1 is sequestered by IGF binding proteins (IGFBPs), which restricts IGF1-IGF1R interactions (Hwa et al., 1999). Given that exogenous IGF1 supplementation can suppress neuron loss (Zheng et al., 2000; Hayashi et al., 2013), the extracellular factors that regulate IGF1 bioavailability may be critical determinants of a neuron’s survival and mitochondrial function. In proximal tubular epithelial cells, IGFBP-3 overexpression has been shown to increase oxidative stress and cell death. Conversely, in these cells knockdown of IGFBP-3 has been shown to suppress toxin-induce oxidative stress and thereby promote cell survival (Yoo et al., 2011). It is currently unknown whether modulation of IGFBPs, and therefore IGF1 bioavailability, also affects neuron survival and mitochondrial function.

To counter the negative regulatory role of IGFBPs, locally secreted proteases cleave IGFBPs to “free” IGF1 and thereby stimulate local IGF1 signaling. One such protease, called Pregnancy-associated plasma protein A (Pappa), has been shown to target a subset of IGFBPs and stimulate multiple IGF1-dependent functions, including synapse formation and function (Boldt and Conover, 2007; Miller et al., 2018). It remains unclear whether Pappa acts as an extracellular regulator of IGF1-dependent neuron survival, mitochondrial function, or oxidative stress. Here, through analysis of a zebrafish *pappaa* mutant, we characterize defects in the survival and mitochondrial function of hair cell sensory neurons and spinal motor neurons. These results reveal a novel role for Pappaa in regulating a neuron’s mitochondrial function and oxidative stress to promote neuron survival.

## Results

### Pappaa regulates survival of hair cell sensory neurons

Zebrafish *pappaa* mutants (hereafter referred to as *pappaa^p170^*) were originally identified based on aberrant behavioral responses to acoustic stimuli (Wolman et al., 2015). In characterizing this phenotype, we assessed the morphology of hair cell sensory neurons, which mediate detection of acoustic stimuli by the inner ear and lateral line sensory organs. Hair cells of the lateral line are clustered into superficially positioned structures called neuromasts (Ghysen and Dambly-Chaudière, 2007) (Fig 1a). At 5 days post fertilization (dpf), *pappaa^p170^* larvae hair cells appeared morphologically indistinguishable from wildtype (Fig 1b). To assess the contribution of hair cell function to the *pappaa^p170^* larvae’s behavioral defects, we briefly exposed 5 dpf *pappaa^p170^* and wild type larvae to neomycin, an aminoglycoside that kills hair cells of the lateral line by disrupting mitochondrial function (Esterberg et al., 2014; Esterberg et al., 2016). 4 hours after neomycin exposure, hair cells of *pappaa^p170^* larvae showed a greater reduction compared to wild type hair cells, suggesting an increased sensitivity to neomycin (Fig 1c-d). Support cells, which surround the hair cell rosettes in each neuromast (Ghysen and Dambly-Chaudière, 2007; Thomas et al., 2015), were unaffected by neomycin exposure (S1a-b Fig). Next, we asked whether Pappaa deficiency yielded naturally occurring hair cell death (i.e. without neomycin treatment). To address this, we evaluated the number of hair cells per neuromast from 5-12 dpf. Within this period, we observed an increase in hair cells in wild type larvae, but not in *pappaa^p170^* larvae (Fig 1e). Because hair cells regenerate in zebrafish (Harris et al., 2003), even in *pappaa^p170^* (S2a Fig), a failure of *pappaa^p170^* hair cells to increase in number suggests their natural degeneration.

**Fig 1.**
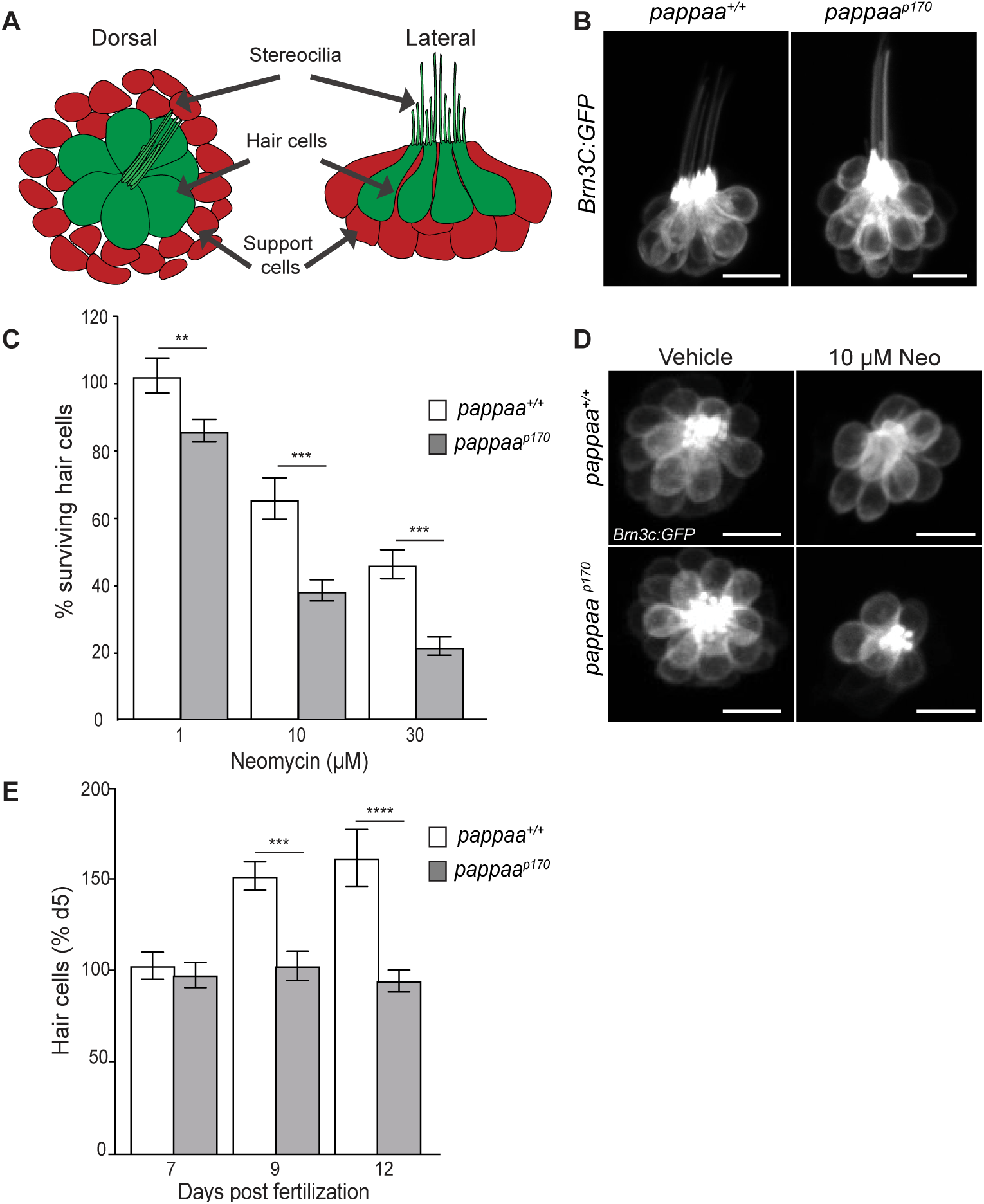
Hair cell survival is reduced in zebrafish *pappaa^p170^* larvae. (a) Schematic of lateral line neuromast. (b) *Brn3c:GFP* labeled hair cells clustered within a lateral line neuromast of wild type and *pappaa^p170^*. Scale = 10µm. (c) Mean percentage of surviving hair cells. To calculate hair cell survival percentage, hair cell number 4 hours post-neomycin treatment was normalized to mean hair cell number in vehicle treated larvae of the same genotype. \*\**p*<0.01, \*\*\**p*<0.001, two-way ANOVA, Holm-Sidak post test. N=13 larvae per group, 3 neuromasts/larva. (d) Representative images of *Brn3c:GFP* labeled hair cells from vehicle or 10µM neomycin treated larvae. Scale = 10µm. (e) Mean percentage of hair cells at 7, 9, and 12 dpf. To calculate hair cell survival percentage, hair cell number at each time point was normalized to mean hair cell number at 5 dpf for a given genotype. \**p*<0.05, \*\*\**p*<0.001, two-way ANOVA, Holm-Sidak post test. N=8 larvae per group, 3 neuromasts/larva. Error bars=SEM.

### Pappaa is expressed by neuromast support cells and motor neurons

To begin to characterize how Pappaa regulates hair cell survival we evaluated *pappaa* mRNA expression in lateral line hair cells and their surrounding environment. *In situ* hybridization revealed *pappaa* expression in lateral line neuromasts with clear expression in the support cells (Fig 2a-b). To determine whether hair cells also express *pappaa* we performed RT-PCR on fluorescently sorted hair cells from 5 dpf *Tg(brn3c: GFP)*(Xiao et al., 2005) larvae (Fig 2c). We did not observe *pappaa* expression by hair cells, suggesting that supports cells express *pappaa* to influence hair cell survival. The *in situ* analysis also indicated *pappaa* expression in the ventral spinal cord, where motor neurons reside (Fig 2a). RT-PCR of fluorescently sorted motor neurons from 5 dpf *Tg(mnx1: GFP)*(Rastegar et al., 2008) larvae confirmed *pappaa* expression by motor neurons (Fig 2c).

**Fig 2.**
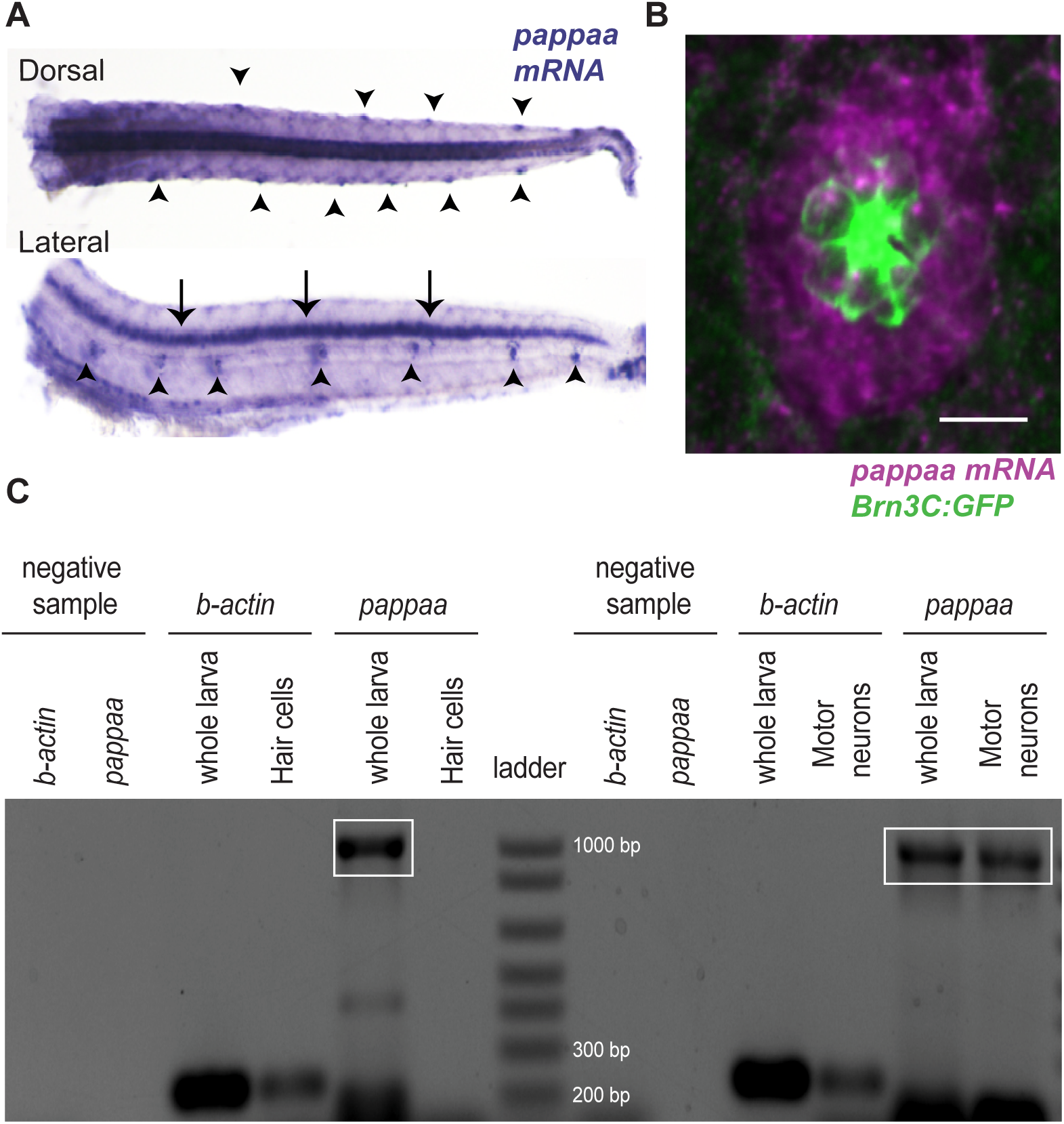
*pappaa* is expressed by neuromast support cells and motor neurons. (a) Whole mount *in situ* hybridization shows *pappaa* mRNA expression at 4 dpf by lateral line neuromasts (arrowheads) and in the ventral spinal cord (arrows). Top image: dorsal view, bottom image: lateral view. (b) Fluorescent in situ hybridization of *pappaa* (magenta) and *Brn3c:GFP* labeled hair cells (green) shows *pappaa* mRNA expression by the support cells that surround hair cells. Scale = 10µm. (c) RT-PCR of fluorescently sorted *Brn3c:GFP* labeled hair cells and *mnx1:GFP* labeled motor neurons shows *pappaa* expression by motor neurons, but not hair cells.

### Pappaa acts through IGF1R signaling to promote neuron survival

A role for Pappaa in neuron survival is novel. Therefore, we sought to characterize the molecular pathway by which Pappaa affects neuron survival. Pappaa is a secreted metalloprotease that cleaves IGFBPs and therefore increases local IGF1 availability and activation of IGF1 receptors (IGF1Rs) (Boldt and Conover, 2007). Neuronal functions of Pappaa, including synapse formation and function, have been shown to be IGF1R signaling dependent (Wolman et al., 2015; Miller et al., 2018). *pappaa^p170^* mutants harbor a nonsense mutation upstream of Pappaa’s proteolytic domain and show reduced IGF1R activation in other neural regions of *pappaa* expression (Wolman et al., 2015; Miller et al., 2018). To determine whether *pappaa^p170^* hair cells show reduced IGF1R activity, we immunolabeled wild type and *pappaa^p170^* larvae for phosphorylated IGF1Rs (pIGF1R) (Chablais and Jazwinska, 2010). In *pappaa^p170^* larvae, we observed a reduction in pIGF1R immunolabeling of *pappaa^p170^* hair cells compared to wild type hair cells (Fig 3a-c).

**Fig 3.**
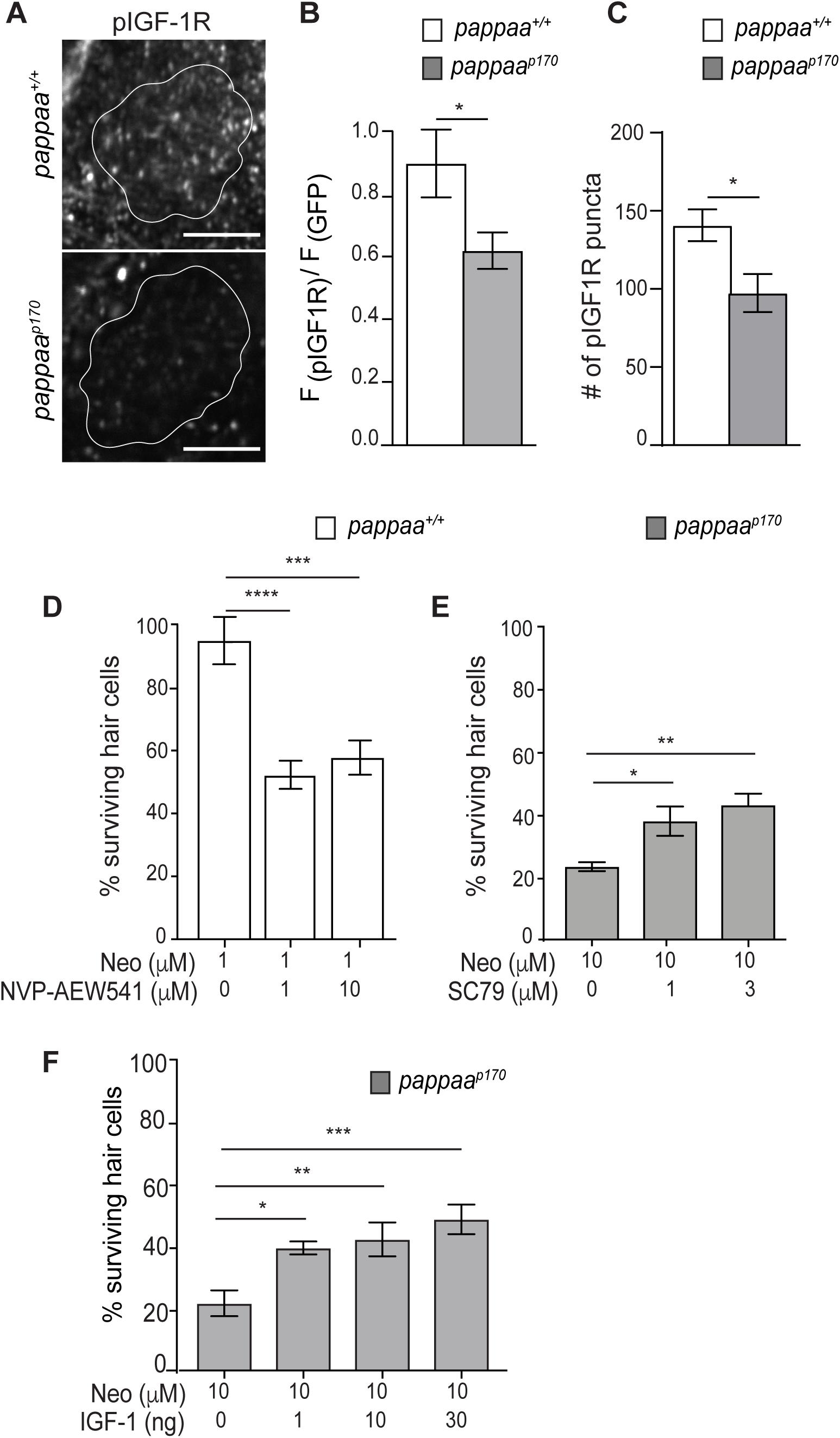
Pappaa - IGF1 signaling regulates hair cell survival. (a) *Brn3c:GFP* marked hair cells (outlined) immunolabeled for phosphorylated IGF1R receptor. Scale = 10µm. (b-c) Mean pIGF1R immunofluorescence normalized to GFP immunofluorescence (b) and mean number of pIGF1R puncta (c) per hair cell cluster. *p<0.05, unpaired *t* test, Welch-corrected. N= 8 larvae, 1-3 neuromasts/larva. (d-f) Mean percentage of surviving hair cells 4 hours post-neomycin treatment. To calculate hair cell survival percentage, hair cell counts in neomycin treated larvae were normalized to vehicle treated larvae of same genotype. (d) Wildtype larvae treated with the IGF1R antagonist NVP-AEW541. (e-f) *pappaa^p170^* larvae treated with SC79 (e) or recombinant IGF1 (f). \**p*<0.05, \*\**p*<0.01, \*\*\**p*<0.001, \*\*\*\**p*<0.0001. One-way ANOVA, Holm-Sidak post test. N=10 larvae, 3 neuromasts/larva. Error bars=SEM.

We next asked whether Pappaa acts via IGF1R signaling to promote hair cell survival. We hypothesized that if Pappaa acts through IGF1R signaling, then attenuating IGF1R activity would reduce hair cell survival following neomycin exposure. To test this hypothesis, we treated wild type larvae with a selective inhibitor of IGF1R phosphorylation, NVP-AEW541 (Chablais and Jazwinska, 2010), for 24 hours prior to and during 1µM neomycin exposure. 1µM neomycin alone did not induce hair cell death (Figs 1c and 3d); however, larvae pre-treated with NVP-AEW541 showed significant hair cell loss after 1µM neomycin exposure (Fig 3d). Next, we hypothesized that if Pappaa acts through IGF1R signaling, then stimulating either IGF1 availability or a downstream effector of the IGF1R would improve hair cell survival in *pappaa^p170^* larvae. To test this hypothesis, we bathed wild type and *pappaa^p170^* larvae in recombinant human IGF1 protein or a small molecule activator of Akt (SC79), a canonical downstream effector of IGF-1R signaling (Laviola et al., 2007). Pre-treatment with IGF1 or SC79 for 24 hours prior to and during neomycin exposure improved hair cell survival in *pappaa^p170^* larvae (Fig 3e-f). Together, these results suggest that Pappaa promotes hair cell survival by stimulating IGF1R signaling.

### Pappaa loss causes increased mitochondrial ROS in hair cells

To define how Pappaa activity influences neuron survival, we evaluated known mechanismsunderlying neomycin-induced hair cell death. Neomycin enters hair cells via mechanotransduction (MET) channels found on the tips of stereocilia (Alharazneh et al., 2011). We hypothesized that *pappaa^p170^* hair cells may be more susceptible to neomycin-induced death due to an increase in MET channel-mediated entry. To assess entry via MET channels we compared uptake of FM1-43, a fluorescent styryl dye that enters cells through MET channels (Meyers et al., 2003). FM1-43 fluorescence was equivalent between wild type and *pappaa^p170^* hair cells (Fig 4a-b), suggesting that the increased death of *pappaa^p170^* hair cells was not due to increased neomycin entry.

**Fig 4.**
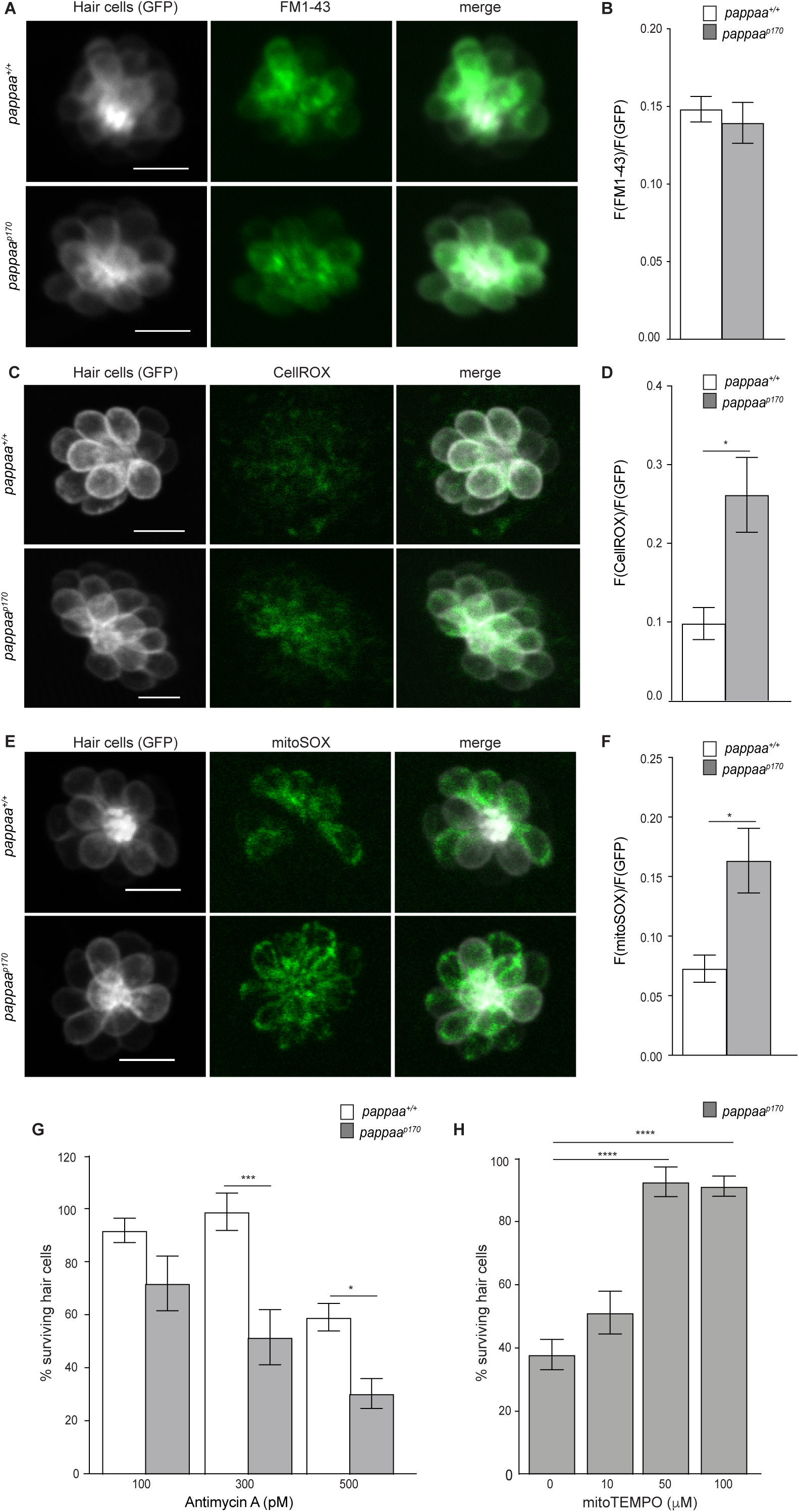
Pappaa regulates mitochondrial ROS generation. (a, c, e) Still images of live *Brn3c:GFP* hair cells loaded with the amphypathic styryl dye FM1-43 (a) or cytoplasmic or mitochondrial ROS indicators (c: CellROX, e: mitoSOX). Scale = 10µm. (b, d, f) Mean dye fluorescence normalized to GFP fluorescence. For b, N= 6 larvae, 2-4 neuromasts/larva. For d, N= 5-6 larvae per group, 2-6 neuromasts/ larva. For f, N= 6 larvae per group, 2-4 neuromasts/ larva. \**p*<0.05. Unpaired *t* test, Welch-corrected. (g) Mean percentage of surviving hair cells post 24 hours incubation in antimycin A. To calculate hair cell survival percentage, hair cell counts after treatment were normalized to hair cell number in vehicle treated larvae of same genotype. \**p*<0.05 \*\*\*\**p*<0.0001. Two-way ANOVA, Holm-Sidak post test. N= 9-10 larvae, 3 neuromasts/larva. (h) Mean percentage of surviving *pappaa^p170^* hair cells following co-treatment with mitoTEMPO and neomycin. To calculate hair cell survival percentage, hair cell counts 4 hours post-neomycin treatment were normalized to hair cell counts in vehicle treated *pappaa^p170^*larvae. \*\*\*\**p*<0.0001. One-way ANOVA, Holm-Sidak post test. N=4-10 larvae, 3 neuromasts/larva. Error bars=SEM.

We therefore hypothesized that Pappaa affects essential organelle functions in hair cells. Within the hair cell, neomycin triggers Ca^2+^ release from the endoplasmic reticulum (ER), which is then taken up by mitochondria (Esterberg et al., 2014). This Ca^2+^ transfer results in stimulation of the mitochondrial respiratory chain, increased mitochondrial transmembrane potential, and an ensuing increase in ROS production (Gorlach et al., 2015; Esterberg et al., 2016). The oxidative stress caused by high ROS levels ultimately underlies the neomycin-induced hair cell death. To explore whether excessive ROS production underlies *pappaa^p170^* hair cells’ increased sensitivity to neomycin, we evaluated cytoplasmic ROS levels with a live fluorescent indicator of ROS (CellROX) (Esterberg et al., 2016). *pappaa^p170^* hair cells displayed elevated ROS levels at baseline; prior to addition of neomycin (Fig 4c-d). Given that the mitochondria are the primary generators of cellular ROS (Lenaz, 2001), we asked whether the elevated levels of cytoplasmic ROS observed in *pappaa^p170^* hair cells originated from the mitochondria. We evaluated mitochondrial ROS with the live fluorescent indicator mitoSOX (Esterberg et al., 2016), again without neomycin treatment, and observed increased signal in hair cells of *pappaa^p170^* compared to wild type (Fig 4e-f). This increased mitochondrial ROS was not due to an overabundance of mitochondria within *pappaa^p170^* hair cells (Fig. S3a-b).

We hypothesized that the elevated ROS in *pappaa^p170^* hair cells predisposed them closer to a cytotoxic threshold of oxidative stress that results in cell death. To test this idea, we asked whether *pappaa^p170^* hair cells were more sensitive to pharmacological stimulation of mitochondrial ROS. To stimulate ROS, we exposed wild type and *pappaa^p170^* larvae to Antimycin A, an inhibitor of the mitochondrial electron transport chain (Hoegger et al., 2008; Quinlan et al., 2011) We found that *pappaa^p170^* hair cells were more susceptible to death by Antimycin A than wild type hair cells (Fig 4g). We next asked whether the increased mitochondria-generated ROS levels in *pappaa^p170^* hair cells underlies their vulnerability to death. We hypothesized that if this were the case, then reducing mitochondrial-ROS would suppress their increased sensitivity to neomycin. To test this idea we exposed *pappaa^p170^* larvae to the mitochondria-targeted ROS scavenger mitoTEMPO (Esterberg et al., 2016) and observed up to complete protection of *pappaa^p170^* hair cells against neomycin-induced death (Fig 4h). These results suggest that abnormally elevated mitochondrial ROS underlies hair cell death in *pappaa^p170^*.

### Pappaa regulates mitochondrial Ca^2+^ uptake and transmembrane potential

Mitochondrial ROS production is stimulated by Ca^2**+**^ entry into the mitochondria (Brookes et al., 2004; Gorlach et al., 2015). Given the increased mitochondrial ROS in *pappaa^p170^* hair cells, we asked whether the mutants’ hair cell mitochondria exhibited increased Ca^2**+**^ levels. To address this, we used a transgenic line *Tg(myo6b:mitoGCaMP3),* in which a mitochondria-targeted genetically encoded Ca^2**+**^ indicator (*GCaMP3*) is expressed in hair cells (Esterberg et al., 2014). Live imaging of mitoGCaMP3 fluorescence revealed a doubling in fluorescent intensity in *pappaa^p170^* hair cells compared to wild type hair cells (Fig 5a-b). Mitochondrial Ca^2**+**^ uptake is driven by the negative electrochemical gradient of the mitochondrial transmembrane potential, a product of mitochondrial respiration. Ca^2**+**^-induced stimulation of mitochondrial oxidative phosphorylation causes further hyperpolarization of mitochondrial transmembrane potential, leading to increased uptake of Ca^2**+**^ (Brookes et al., 2004; Adam-Vizi and Starkov, 2010; Ivannikov and Macleod, 2013; Esterberg et al., 2014; Gorlach et al., 2015). Therefore, we hypothesized that *pappaa^p170^* mitochondria would have a more negative transmembrane potential compared to wild type. Using the potentiometric probe TMRE that provides a fluorescent readout of mitochondrial transmembrane potential (Perry et al., 2011), we found that *pappaa^p170^* mitochondria possess a more negative transmembrane potential compared to wild type (Fig 5c-d). This result is consistent with the *pappaa^p170^* mitochondria’s increased Ca^2+^ load. Given that mitochondria of *pappaa^p170^* hair cells exhibit elevated Ca^2**+**^ (Fig 5a-b) and a more negative transmembrane potential (Fig 5c-d) at baseline, we hypothesized that pharmacologically disrupting these mitochondrial features would have a more cytotoxic effect on *pappaa^p170^* hair cells. To test this idea, we exposed wild type and *pappaa^p170^* larvae to Cyclosporin A (CsA), an inhibitor of the mitochondrial permeability transition pore that causes buildup of mitochondrial Ca^2+^ and further hyperpolarizes mitochondria (Crompton et al., 1988; Esterberg et al., 2014). *pappaa^p170^* larvae showed reduced hair cell survival at concentrations of CsA, which had no effect on hair cell survival in wild type larvae (Fig 5e). Taken together, these results suggest that Pappaa regulates mitochondrial ROS production by attenuating mitochondrial Ca^2**+**^ uptake.

**Fig 5.**
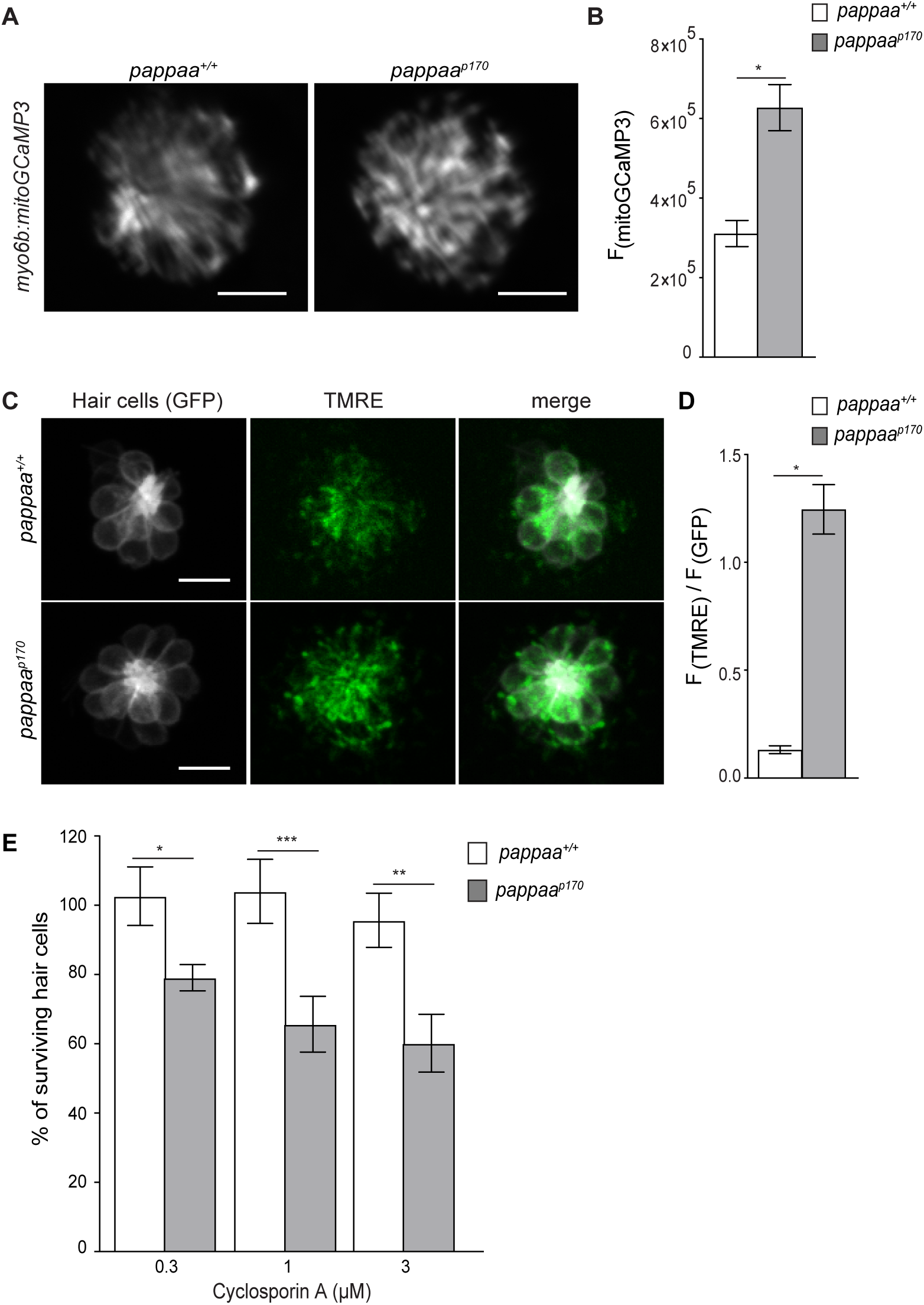
Mitochondrial Ca^2+^ levels and transmembrane potential are disrupted in *pappaa^p170^* hair cells. (a) Still images from live *myo6b:mitoGCaMP3* labeled hair cells. Scale = 10µm. (b) Mean mitoGCaM*P* fluorescence. \**p*<0.05. Unpaired *t* test, Welch-corrected. N= 4-6 larvae, 3-4 neuromasts/larva. (c) Still images from live *Brn3c:GFP* labeled hair cells loaded with TMRE. Scale = 10µm. (d) Mean TMRE fluorescence normalized to GFP fluorescence. N= 4 larvae, 3-4 neuromasts/larva. \**p*<0.05. Unpaired *t* test, Welch-corrected. (e) Mean percentage of surviving hair cells post Cyclosporin A treatment. To calculate hair cell survival percentage, hair cell counts post-treatment were normalized to hair cell numbers in vehicle treated larvae of same genotype. \**p*<0.05, \*\**p*<0.01, \*\*\**p*<0.001. Two-way ANOVA, Holm-Sidak post test. N=8-13 larvae, 3 neuromasts/larva. Error bars=SEM.

### Pappaa deficient motor neurons show degeneration and oxidative stress

We were curious to know whether Pappaa promotes the survival of other neuron types by attenuating oxidative stress. To address this, we evaluated the number of spinal motor neurons in wild type and *pappaa^p170^* larvae at 5 and 9 dpf. Using the *Tg(mnx1:GFP)* line to count motor neurons, we observed a reduced number of motor neurons and thinning of the ventral projecting motor nerve in 9 dpf *pappaa^p170^* larvae (Fig 6a-c). To determine whether *pappaa^p170^* motor neurons also exhibit oxidative stress we analyzed antioxidant gene expression in motor neurons by RT-qPCR. An increase in antioxidant gene expression is an adaptive response to elevated ROS (Gorrini et al., 2013; Syu et al., 2016). qPCR of cDNA from fluorescently sorted *Tg(mnx1:GFP)* motor neurons revealed increased expression of several antioxidants genes (*gpx*, *sod2*, and *catalase*) in *pappaa^p170^* compared to wild type (Fig 6d). Taken together, these results reveal that Pappaa’s influence on oxidative stress and survival affects multiple neuron types.

**Fig 6.**
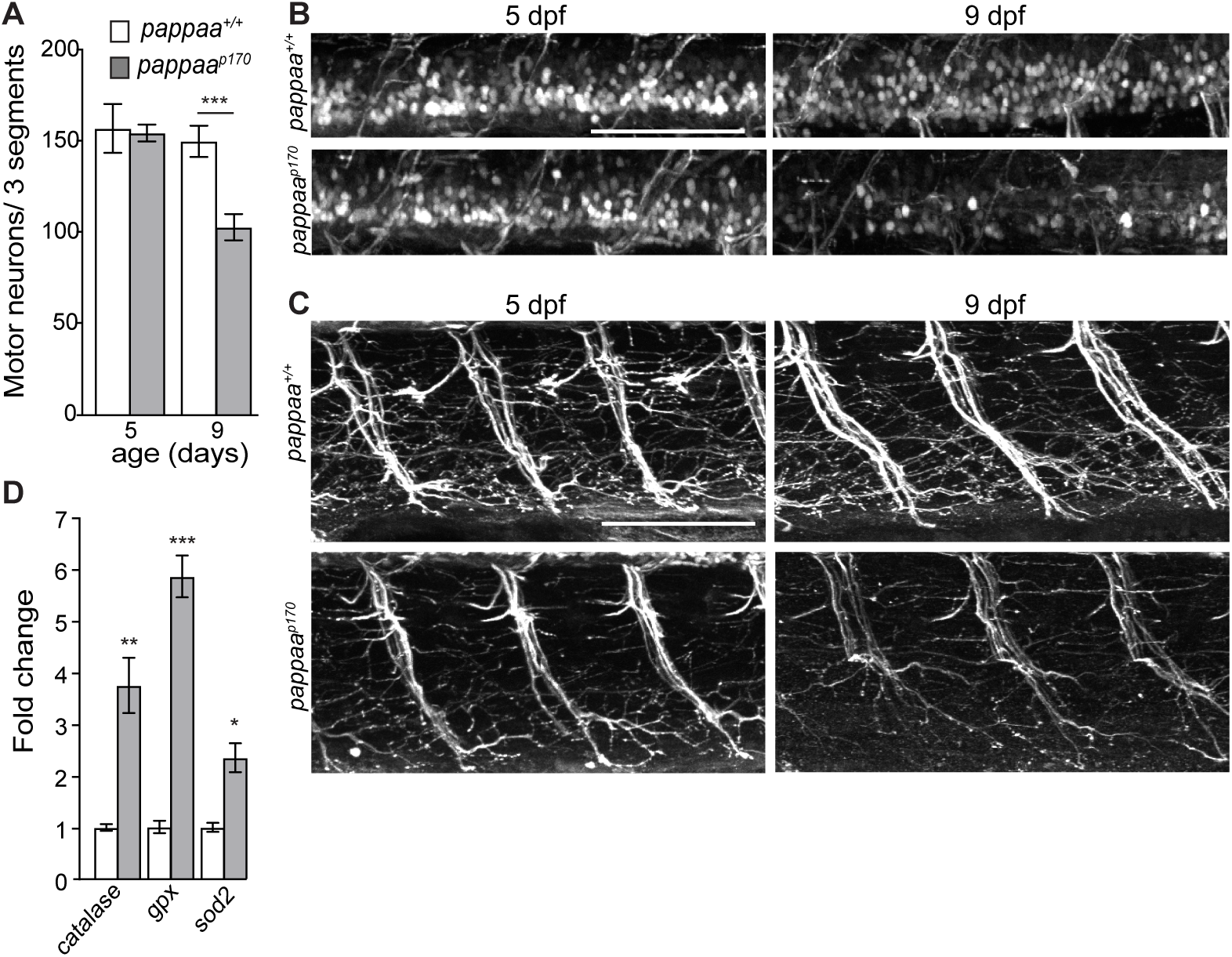
Pappaa-deficient spinal motor neurons die precociously. (a) Mean number of *mnx1:GFP* labeled spinal motor neurons summed across 3 motor segments at 5 and 9 dpf. \*\**p*<0.01. Unpaired t-test, Welch corrected. N=6-8 larvae, 3 segments/ larva. (b-c) Representative images of *mnx1:GFP* labeled motor neurons (b) and nerves (c), captured with the spinal cord (c) and innervating the ventral myotome (c) of the same larvae. Scale = 100µm. (d) Mean fold change in antioxidants transcript expression levels in motor neurons at 5dpf. N= 3 technical replicates/gene. \**p*<0.05, \*\**p*<0.01, \*\*\**p*<0.001. Unpaired *t* test, Welch-corrected. Error bars=SEM.

## Discussion

The precise regulation of mitochondrial function and ROS production is essential for neuron survival. Indeed, mitochondrial dysfunction, and the ensuing overproduction of ROS is causative of neuron death (Lin and Beal, 2006). Extracellular molecular factors produced by neurons or non-neuronal cells are known to influence neuron survival (Hasan et al., 2003; Li et al., 2009; Wang et al., 2013; Genis et al., 2014). Yet it remains poorly understood which of these factors support neuron survival by affecting mitochondrial function and ROS production. Here, through zebrafish mutant analysis, we reveal a novel extracellular regulator of neuron survival and mitochondrial function: the secreted metalloprotease Pappaa. Based on a series of *in vivo* experiments we propose a model by which Pappaa stimulates IGF1 receptor signaling in neurons to control mitochondrial function and ROS production, and thereby, promote neuron survival.

### Pappaa-IGF1 receptor signaling is required for neuron survival

Pappaa’s requirement for neuron survival is demonstrated by a precocious loss of sensory hair cells and spinal motor neurons in *pappaa^p170^* zebrafish larvae. Neuron loss was spontaneous for both populations (Figs 1e and 6a-b), and the hair cell loss was accentuated by exposure to mitochondrial toxins (Figs 4g and 5e). Pappaa’s role in neuron survival is novel and therefore it is interesting to consider whether Pappaa serves this role by promoting neuronal development, maintenance, and/or regeneration. In the *pappaa^p170^* mutants, both hair cells and motor neurons appeared to develop normally, based on their numbers and cellular morphology in 5 dpf larvae. At 5 dpf, larvae require these neurons for various behaviors, including eliciting an acoustic startle response (Bang et al., 2002; Wolman et al., 2015). Although *pappaa^p170^* mutant larvae show deficits in startle modulation, we have previously shown that at 5 dpf the mutants have the ability to detect acoustic stimuli and perform explosive escape maneuvers (Wolman et al., 2015). This ability further suggests that *pappaa^p170^* mutant hair cells and motor neurons were functionally intact prior to the onset of their loss, and therefore developed normally. In zebrafish, both hair cells and spinal motor neurons are capable of regeneration (Thomas et al., 2015; Ohnmacht et al., 2016). Therefore, albeit unprecedented, it is possible that these neurons naturally die at the rate we observed in *pappaa^p170^* mutants, but then require Pappaa to regenerate. This possibility is unlikely given our observation that hair cells in *pappaa^p170^* mutants showed a normal regenerative capacity after neomycin-induced loss (S2 Fig). Taken together, these results suggest that Pappaa is dispensable for the development and regeneration of hair cells and spinal motor neurons, and rather, supports their maintenance.

Increased IGF1 signaling has been shown to provide neuroprotection(Zheng et al., 2000). For example, exogenously supplied IGF1 was recently demonstrated to protect hair cells from neomycin-induced damage (Hayashi et al., 2013). However, it is poorly understood how endogenous IGF1 signaling is regulated to promote neuron survival, particularly through extracellular factors. Pappaa is a secreted metalloprotease that cleaves the inhibitory IGF binding proteins, thereby freeing IGF-1 to bind and activate cell-surface IGF1 receptors. Thus, Pappaa acts as an extracellular positive regulator of IGF-1 signaling (Boldt and Conover, 2007). Consistent with this role for Pappaa, immunolabeling of activated, phosphorylated IGF1Rs was reduced on hair cells of 5 dpf *pappaa^p170^* mutants (Fig 3a-c). Stimulation of IGF1R signaling, either by supplementation of IGF1 or by stimulation of the IGF1R effector Akt, suppressed hair cell loss in *pappaa^p170^* mutants (Fig 3e-f). Notably, stimulation of IGF1R signaling after the hair cells had developed was sufficient to suppress this loss, which is consistent with a post-developmental role for Pappaa in regulating hair cell survival.

For hair cells and motor neurons, *pappaa* expression suggests that Pappaa can act in a paracrine or autocrine manner, respectively, to promote neuron survival. Although hair cells require Pappaa for survival, they do not express Pappaa. Rather, their surrounding support cells, expressed *pappaa* (Fig 2a-c). These support cells have been demonstrated to secrete factors that promote hair cell survival (May et al., 2013) and our results suggest that Pappaa is one such factor. In contrast to hair cells, *pappaa* is expressed by the spinal motor neurons that require Pappaa for survival (Figs 2a, 2c, and 6a-b), suggesting an autocrine function. Our analysis has not excluded that neighboring spinal neurons might also provide a Pappaa source for motor neurons. It remains unclear whether motor neuron-secreted Pappaa also promotes the survival of other spinal neuron types. We speculate that it does given the relative ubiquity of neuronal IGF1R expression and the need for all spinal neurons to regulate mitochondrial activity. To understand Pappaa’s cell autonomy it will be necessary to define what triggers Pappaa activity to promote neuron survival. Is Pappaa acting in response to cues from dying neurons, and what are these cues, or is Pappaa serving a preventative role?

### Pappaa regulates mitochondrial function in neurons

The mitochondria in *pappaa^p170^* mutant hair cells showed multiple signs of dysfunction, including elevated ROS (Fig 4c-f), transmembrane potential (Fig 5c-d), and Ca^2**+**^ load (Fig 5a-b). Consistent with these observations, reduced IGF1 signaling has been associated with increased ROS production and oxidative stress (García-Fernández et al., 2008; Lyons et al., 2017). Two lines of evidence suggest that mitochondrial dysfunction, and particularly the elevated ROS production, underlie neuron loss in *pappaa^p170^* mutants. First, *pappaa^p170^* hair cells showed enhanced sensitivity to pharmacological stimulators of mitochondrial ROS production (Figs 4g and 5e). Second, attenuation of mitochondrial ROS by mitoTEMPO exposure, a mitochondrial targeted antioxidant, was sufficient to suppress neomycin-induced hair cell loss in *pappaa^p170^* mutants (Fig 4h).

Based on results presented here, it is difficult to pinpoint the exact locus of mitochondrial dysfunction in *pappaa^p170^* neurons due to the tight interplay between mitochondrial transmembrane potential, Ca^2**+**^ load, and ROS production (Brookes et al., 2004; Adam-Vizi and Starkov, 2010; Ivannikov and Macleod, 2013; Esterberg et al., 2014; Gorlach et al., 2015). The oxidative phosphorylation process that generates ROS relies on maintaining a negative mitochondrial transmembrane potential. Negative transmembrane potential is achieved by pumping protons out of the mitochondrial matrix as electrons move across the electron transport chain. Protons then move down the electrochemical gradient through ATP synthase to produce ATP. Given that ROS is a byproduct of oxidative phosphorylation, a more negative transmembrane potential yields more ROS (Kann and Kovács, 2007; Zorov et al., 2014). Mitochondrial Ca^2+^ is a key regulator of transmembrane potential and the resultant ROS generation, as it stimulates the activity of key enzymes involved in oxidative phosphorylation (Brookes et al., 2004). And, Ca^2**+**^ uptake by the mitochondria is driven by the electrochemical gradient of a negative transmembrane potential. Thus, Ca^2**+**^ and transmembrane potential are locked in a feedback loop (Brookes et al., 2004; Adam-Vizi and Starkov, 2010; Ivannikov and Macleod, 2013; Esterberg et al., 2014; Gorlach et al., 2015). Because mitochondria in *pappaa^p170^* hair cells have a more negative transmembrane potential (Fig 5c-d) and experience Ca^2+^ overload (Fig 5a-b), this likely sensitizes the mitochondria to any further increase in Ca^2+^ levels. In support of this, *pappaa^p170^* hair cells were hypersensitive to Cyclosporin A (Fig 5e), which increases mitochondrial Ca^2+^ levels by blocking the mitochondrial permeability transition pore (Smaili and Russell, 1999).

It is possible that the *pappaa^p170^* neurons dysfunctional mitochondria are downstream effects of anomalies in cellular mechanisms acting outside of the mitochondria. A potential driver of the excessive mitochondrial Ca^2+^ levels and ROS production in *pappaa^p170^* hair cells is the endoplasmic reticulum (ER). ER and mitochondria are structurally coupled to facilitate rapid and efficient transfer of Ca^2**+**^ (Esterberg et al., 2014; Krols et al., 2016). Neomycin, to which the *pappaa^p170^* neurons showed hypersensitivity, stimulates this transfer (Esterberg et al., 2014). ER-mitochondria Ca^2**+**^ transfer can be enhanced by ER stress (Bravo et al., 2012). A cause of ER stress is disruption to ER-mediated protein processing mechanisms. In addition to playing a key role in buffering neuronal Ca^2+^, the ER lumen is a major site for protein processing, including protein folding. Nascent proteins enter the ER to be folded with the aid of molecular chaperones. Insufficient folding yields an accumulation of misfolded proteins, which triggers efflux of Ca^2**+**^ (Deniaud et al., 2008; Houck et al., 2012). IGF1 signaling has been shown to promote the ER’s protein folding capacity, and thereby attenuate ER stress (Barati et al., 2006; Novosyadlyy et al., 2008; Chatterjee et al., 2013). Notably, this relationship between the ER and mitochondria that governs Ca^2**+**^ transfer is not unidirectional. Mitochondria-generated ROS can modulate the activity of ER Ca^2**+**^ channels and cause ER Ca^2**+**^ efflux that further stimulates mitochondrial ROS production (Peng and Jou, 2010; Gorlach et al., 2015). Given that many ER-resident molecular chaperones are Ca^2**+**^-dependent (Gidalevitz et al., 2013), protein misfolding can be both a cause and a consequence of ER-Ca^2**+**^ depletion. Further experimental dissections of ER mediated functions and the interactions between the ER and mitochondria in *pappaa^p17^*mutant neurons are needed to define the primary locus within neurons by which Pappaa-IGF1 signaling influences neuron survival.

Here, we define a novel role for Pappaa in neuron survival by stimulating the IGF1 signaling pathway and regulating mitochondrial function. The evidence we present is consistent with demonstrations that IGF1 regulates neuron survival and mitochondrial function (Zheng et al., 2000; Luo et al., 2003; García-Fernández et al., 2008; Lyons et al., 2017). Our discovery of Pappaa in this context breathes hope into the potential for IGF1 mediated therapies for neurodegenerative diseases. Unfortunately, patients with neurodegenerative disorders have not shown significant symptomatic improvement following systemic IGF1 administration. These disappointing outcomes are thought to be due to the suppressive effects of IGFBPs on IGF1 bioavailability (Raoul and Aebischer, 2004; Sakowski et al., 2009). In support of this, patients with ALS exhibit normal total levels of IGF1, while free IGF1 was reduced likely due to the upregulation of IGFBPs (Wilczak et al., 2003). Furthermore, the ubiquitous nature and broad cellular impact of IGF1 signaling presents a major challenge for a therapy based on enhancing systemic IGF1 (Joseph D'Ercole and Ye, 2008). Temporal and spatial restrictions to IGF1 signaling may yield better outcomes. As a local, upstream regulator of IGF1 signaling with restricted spatial expression, Pappaa may be a viable target to locally stimulate IGF1 signaling to combat neuron loss in disease.

## Materials and Methods

### Maintenance of Zebrafish

To generate *pappaa^+/+^* and *pappaa^p170^* larvae for experimentation, adult *pappaa^p170/+^* zebrafish (on a mixed Tubingen long-fin, WIK background) were crossed into the following transgenic zebrafish backgrounds: *Tg(brn3c:GFP)^s356t^, Tg(mnx1:GFP)^mI5^,* and *Tg(myo6b:mitoGCaMP3)^w78^* and then incrossed. Embryonic and larval zebrafish were raised in E3 media (5 mM NaCl, 0.17 mM KCl, 0.33 mM CaCl_2_, 0.33 mM MgSO_4_, pH adjusted to 6.8–6.9 with NaHCO_3_) at 29°C on a 14 hour/10 hour light/dark cycle through 5 days post fertilization (dpf) (Kimmel et al., 1995; Gyda et al., 2012). Larvae raised beyond 5 dpf were fed paramecia. All experiments were done on 4-12 dpf larvae. Genotyping of *pappaa^p170^* larvae was performed as previously described (Wolman et al., 2015).

### Pharmacology

The following treatments were performed on *Tg(brn3c:GFP)* larvae through the addition of compounds to the larvae’s E3 media at 5 dpf unless otherwise noted. Neomycin sulfate solution (Sigma-Aldrich) was added at 1-30 μM for 1 hour. Cyclosporin A (Abcam; dissolved in DMSO) was added at 0.3-3 μM for 1 hour. Antimycin A (Sigma-Aldrich; dissolved in DMSO) was added at 100- 500 pM for 24 hours, beginning at 4 dpf. MitoTEMPO (Sigma-Aldrich; dissolved in DMSO) was added at 10-100 μM 30 minutes prior to a 1-hour exposure to 10 μM neomycin. To modulate IGF1 signaling: larvae were pre-treated with NVP-AEW541 at 1-10 μM (Selleck; dissolved in DMSO), SC79 at 1-3 μM (Tocris Bioscience, dissolved in DMSO), or recombinant IGF1 at 1-30 ng/mL (Cell Sciences; dissolved in 10 μM HCl) for 24 hours prior (beginning at 4 dpf) and then exposed to 1-10 μM neomycin for 1 hour on 5 dpf. Following each treatment period, larvae were washed 3 times with E3 and left to recover in E3 for 4 hours at 28°C before fixation with 4% paraformaldehyde (diluted to 4% w/v in PBS from 16% w/v in 0.1M phosphate buffer, pH 7.4). For mitoTEMPO, NVP-AEW541, SC79, and IGF1 treatment, the compounds were re-added to the E3 media for the 4-hour recovery period post neomycin washout. Vehicle-treated controls were exposed to either 0.9% sodium chloride in E3 (neomycin control), E3 only (IGF1 control), or 1% DMSO in E3 for the remaining compounds.

### Hair cell survival

Hair cell survival experiments were performed in *Tg(brn3c:GFP)* larvae where hair cells are marked by GFP. For each larva, hair cells were counted from the same 3 stereotypically positioned neuromasts (IO3, M2, and OP1) (Raible and Kruse, 2000) and averaged. The percent of surviving hair cells was calculated as: [(mean number of hair cells after treatment)/ (mean number of hair cells in vehicle treated group)] X 100. For analyses of neuron survival over time, we normalized the number of neurons counted at each timepoint to the number of neurons present at 5 dpf. Normalizations were genotype specific to account for a slight increase in hair cell number (~2 per neuromast) in *pappaa^p170^* larvae at 5 dpf.

### Single cell dissociation and fluorescence activated cell sorting

For each genotype, 30 5 dpf *Tg(mnx1:GFP)* and 200 5 dpf *Tg(brn3c:GFP)* larvae were rinsed for 15 minutes in Ringer’s solution (116mM NaCl, 2.9 mM KCl, 1.8 mM CaCl_2_, 5 mM HEPES, pH 7.2)(Guille, 1999). *pappaa^p170^* larvae were sorted by swim bladder(Wolman et al., 2015). To collect motor neurons we used whole *Tg(mnx1:GFP)* larvae and to collect hair cells we used tails dissected from *Tg(brn3c:GFP)* larvae. Samples were pooled into 1.5 mL tubes containing Ringer’s solution on ice, which was then replaced with 1.3 mL of 0.25% trypsin-EDTA for digestion. *Tg(mnx1:GFP)* samples were incubated for 90 minutes and *Tg(brn3c:GFP)* were incubated for 20 minutes. Samples were titrated gently by P1000 pipette tip every 15 minutes for motor neurons and every 5 minutes for hair cells. To stop cell digestion, 200µL of 30% FBS and 6 mM CaCl_2_ in PBS solution (Steiner et al., 2014) was added, cells were centrifuged at 400g for 5 minutes at 4°C, the supernatant was removed, the cell pellet was rinsed with Ca^2+^-free Ringer’s solution and centrifuged again. The cell pellet was then resuspended in 1X Ca^2+^-free Ringer’s solution (116mM NaCl, 2.9 mM KCl, 5 mM HEPES, pH 7.2) and kept on ice until sorting. Immediately before sorting, cells were filtered through a 40 µm cell strainer and stained with DAPI. A two-gates sorting strategy was employed. DAPI was used to isolate live cells, followed by a forward scatter (FSC) and GFP gate to isolate GFP+ cells. Sorted cells were collected into RNAse-free tubes containing 500 µL of TRIzol reagent (Invitrogen) for RNA extraction.

### RNA extraction and RT-PCR

Total RNA was extracted from whole larvae and FACS sorted motor neurons and hair cells using TRIzol. cDNA was synthesized using SuperScript II Reverse Transcriptase (Invitrogen). Real-time Quantitative PCR (RT-qPCR) was performed using Sso fast Eva Green Supermix (Biorad) in a StepOnePlus Real-Time PCR System (Applied Biosystems) based on manufacture recommendation. Reactions were run in triplicates containing cDNA from 50 ng of total RNA/reaction. The primer sequences for the antioxidant genes were previously described(Jin et al., 2010) and are as follows: For *sod1,* forward: GTCGTCTGGCTTGTGGAGTG and reverse: TGTCAGCGGGCTAGTGCTT; for *gpx,* forward: AGATGTCATTCCTGCACACG and reverse: AAGGAGAAGCTTCCTCAGCC; for *catalase,* forward: AGGGCAACTGGGATCTTACA and reverse: TTTATGGGACCAGACCTTGG. *b-actin* was used as an endogenous control with the following primer sequences: forward TACAGCTTCACCACCACAGC and reverse: AAGGAAGGCTGGAAGAGAGC(Wang et al., 2005). Cycling conditions were as follows: 1 min at 95°C, then 40 cycles of 15 sec at 95°C, followed by 1 min at 60°C (Jin et al., 2010): Relative quantification of gene expression was done using the 2^−ΔΔCt^method (Livak and Schmittgen, 2001). PCR amplification for *pappaa* fragment was performed by using forward primer: AGACAGGGATGTGGAGTACG, and reverse primer: GTTGCAGACGACAGTACAGC. PCR conditions were as follows: 3 min at 94°C, followed by 40 cycles of 94°C for 30 sec, 57°C for 1 min, and 70°C for 1 min^(Wolman et al., 2015)^. The PCR product was run on a 3% agarose gel.

### Live imaging

All experiments were done on 5-6 dpf *pappaa^p170^* and *pappaa^+/+^* larvae at room temperature. Images were acquired with an Olympus Fluoview confocal laser scanning microscope (FV1000) using Fluoview software (FV10-ASW 4.2). To detect oxidative stress, *Tg(brn3c:GFP)* larvae were incubated in 10 μM CellROX Deep Red (Thermofischer Scientific C10422; dissolved in DMSO) and 1 μM mitoSOX Red (Thermofischer Scientific M36008; dissolved in DMSO) in E3 for 60 minutes and 30 minutes, respectively. To detect transmembrane potential, *Tg(brn3c:GFP)* larvae were incubated in 25 nM TMRE (Thermofischer Scientific T669; dissolved in DMSO) for 20 minutes. To detect MET channel function, *Tg(brn3c:GFP)* larvae were incubated in 3 μM FM1-43 (Thermofischer Scientific T3136; dissolved in DMSO) for 30 seconds. To measure mitochondrial mass, larvae were incubated in 100 nM mitotracker green FM (Thermofischer Scientific M7514; dissolved in DMSO) for 5 minutes. Following the incubation period, larvae were washed 3 times in E3, anesthetized in 0.002% tricaine (Sigma-Aldrich) in E3, and mounted as previously described(Stawicki et al., 2014). Fluorescent intensity of the reporter was measured using ImageJ by drawing regions of interest around hair cells of the neuromast from Z-stack summations. The corrected total cell fluorescence (CTCF) of each reporter was calculated using the following formula: Integrated Density - (Area of selected cells X Mean fluorescence of background) (McCloy et al., 2014). Relative fluorescent intensity was reported as the ratio to GFP fluorescence.

### Immunohistochemistry and in situ hybridization

For whole-mount immunostaining, larvae at 5 dpf were fixed in 4% paraformaldehyde for 1 hour at room temperature. *Tg(mnx1:GFP)* larvae were permeabilized in collagenase (0.1% in PBS) for 4 hours. Larvae were blocked for 1 hour at room temperature in incubation buffer (0.2% bovine serum albumin, 2% normal goat serum, 0.8% Triton-X, 1% DMSO, in PBS, pH 7.4). Larvae were incubated in primary antibodies in IB overnight at 4°C. Primary antibodies were as follows: phosphorylated IGF1R (anti-IGF1 receptor phospho Y1161, 1:100, rabbit IgG; Abcam), hair cells using *Tg(brn3c:GFP)* larvae (anti-GFP, 1:500, rabbit IgG; ThermoFisher Scientific), motor neurons using *Tg(mnx1:GFP)* larvae (anti-GFP, 1:500, rabbit IgG), and support cells (anti-SOX2 ab97959, 1:200, rabbit IgG; Abcam)(He et al., 2014). Following incubation of primary antibodies, larvae were incubated in fluorescently conjugated secondary antibodies in IB for 4 hours at room temperature. Secondary antibodies included AlexaFluor488-conjugated and AlexaFluor594-conjugated secondary antibodies (goat anti-mouse IgG and IgG1, goat anti-rabbit IgG, 1:500; ThermoFisher Scientific). After staining, larvae were mounted in 70% glycerol in PBS. Images were acquired with an Olympus Fluoview confocal laser scanning microscope (FV1000) using Fluoview software (FV10-ASW 4.2).

For whole-mount in situ hybridization: digoxygenin-UTP-labeled antisense riboprobes for *pappaa* (Wolman et al., 2015) were used as previously described (Halloran et al., 1999; Chalasani et al., 2007). Images of colorimetric *in situ* reactions were acquired using a Leica Fluorescence stereo microscope with a Leica DFC310 FX digital color camera. Images of fluorescent *in situ* reactions were acquired using an Olympus Fluoview confocal laser scanning microscope (FV1000).

### Statistics

All data were analyzed using GraphPad Prism Software 7.0b (GraphPad Software Incorporated, La Jolla, Ca, USA). Prior to use of parametric statistics, the assumption of normality was tested using Brown-Forsythe test and Bartlett’s test. Parametric analyses were performed using an unpaired T-test with Welch’s correction or ANOVAs with a Holm-Sidak correction. Data are presented as means ± standard error of the mean (SEM; n = sample size). Significance was set at p < 0.05. N for each experiment is detailed in the results and/or figure legends.

## Author details

### Mroj Alassaf

- Department of Integrative Biology. University of Wisconsin, Madison, Wisconsin, United States of America.
- Neuroscience Training Program. University of Wisconsin, Madison, Wisconsin, United States of America.

### Contribution

Conceptualization, Investigation, Data curation, Formal analysis, Writing – original draft preparation.

### Emily Daykin

Department of Integrative Biology. University of Wisconsin, Madison, Wisconsin, United States of America.

### Contribution

Data curation

### Marc Wolman

Department of Integrative Biology. University of Wisconsin, Madison, Wisconsin, United States of America.

### Contribution

Conceptualization, Resources, Writing – review & editing, Supervision, Project administration, Funding acquisition.

### Funding

### Ministry of Education-Saudi Arabia

- Mroj Alassaf

### University of Wisconsin Sophomore research fellowship and the College of Agricultural & Life Sciences summer undergraduate research award

- Emily Daykin

### Greater Milwaukee Foundation Shaw Scientist Award (133-AAA265)

- Marc Wolman

The funders had no role in study design, data collection and interpretation, or the decision to submit the work for publication

## Acknowledgments

The authors would like to thank Dr. David Raible (University of Washington-Seattle) for the *myo6b:mitoGCaMP3* fish line and Dr. Corinna Burger (University of Wisconsin Department of Neurology) for use of the RT-qPCR cycler.

## Supporting information

**S1 Fig. Support cells are not affected by neomycin treatment.** (a) Mean percentage of surviving neuromast support cells at 4 hours post-neomycin treatment. To calculate support cell survival percentage, support cell number 4 hours post-neomycin treatment was normalized to mean support cell number in vehicle treated larvae in the same genotype. Two-way ANOVA, Holm-Sidak post test revealed no significant difference among groups. N= 6-8 larvae per group, 3-4 neuromasts/ larva. Error bars = SEM. (b) Representative confocal images of *anti-SOX2* labeled support cells that were control or 30µM neomycin treated. Scale = 10µm.

**S2 Fig. *pappaa^p170^* hair cells regenerate and display normal gross morphology.** Mean percentage of hair cells in *pappaa^p170^* at 4, 24, and 48 hours following neomycin treatment at 5 dpf. To calculate hair cell percentage, hair cell number at each timepoint was normalized to mean hair cell number in vehicle treated larvae at 5 dpf. Error bars=SEM.

**S3 Fig. *pappaa^p170^* hair cells have normal mitochondrial mass.** (a) Still images from live wild type and *pappaa^p170^* hair cells loaded with the mitochondrial mass marker, Mitotracker. Scale = 10µm. (b) Mean mitotracker fluorescence. Unpaired *t* test with Welch correction revealed no significant difference among groups. N=3-5 neuromasts, 3-4 neuromasts/larva. Error bars=SEM.

